# Partner-assisted artificial selection of a secondary function for efficient bioremediation

**DOI:** 10.1101/2022.05.12.491680

**Authors:** Marco Zaccaria, Natalie Sandlin, Yoav Soen, Babak Momeni

## Abstract

Microbial enzymes have a broad potential to address many current needs, such as detoxification of harmful toxins and waste, but their native performance often does not match specific applications of interest. In attempting to evolve strains for a specific need, one challenge is that our functions of interest may not confer a fitness effect on the producer. As a result, a conventional selection scheme cannot be used to improve such secondary functions. We propose an alternative approach, partner-assisted artificial selection (PAAS), in which an assisting population acts as an intermediate to create a feedback from the function of interest to the fitness of the producer. We use a simplified model to examine how well and under what conditions such a scheme leads to improved enzymatic function, focusing on degradation of a toxin as a case example. We find that selection for improved growth in this scheme successfully leads to improved degradation performance, even in the presence of other sources of stochasticity. We find that standard selection considerations apply in PAAS: a more restrictive bottleneck leads to stronger selection but adds uncertainty. We also examine how much stochasticity in other traits can be tolerated in PAAS. Our findings offer a roadmap for successful implementation of PAAS to evolve improved functions of interest such as detoxification of harmful compounds.

## Introduction

The vast diversity of bacterial and fungal enzymes offers potential solutions to many current challenges, including the removal of toxic compounds. Recycling complex compounds is an integrated part of the life-style for many bacteria and fungi. The same enzymes can potentially target and remove toxins that contaminate our food, water, and environment. One hurdle in employing native bacterial and fungal enzymes is that the function they are adapted for may not match the degradation of our toxins of interest. As a result, the degradation performance will not meet the demands for practical applications. How can we improve such enzymatic functions? Selection for improved activity would be a clear choice, but what if enzymatic activity against such toxins is a secondary function, where toxin presence or degradation has no direct fitness impact on the bacterial or fungal cells that produce the enzyme?

An illustrative example is the bacterial degradation of mycotoxins—fungal produced food contaminants that are toxic to consume. There are several bacteria and fungi that have already been identified to carry enzymes that degrade mycotoxins [1–6]. However, at least in some cases, the presence of the toxin has little impact on the growth of bacterial cells, posing a challenge for selection. To show an example of such a situation, we have tested the growth of *Rhodococcus erythropolis* cells under different concentrations of aflatoxin G2 (AFG_2_) in the culture (Fig S1). Even though *R. erythropolis* is known to degrade aflatoxins [7–9], AFG_2_ has little positive or negative impact on its growth.

To implement a selection scheme for improving secondary microbial functions, such as detoxification of AFG_2_ by *Rhodococcus*, the detoxification performance should be linked to the detoxifier’s fitness. We propose adding an “assisting” partner population that provides the fitness feedback from the toxin to the detoxifier (Fig 1). Community evolution has recently been revisited for its potential to improve community functions [10–12]. Here we take a slightly different approach by designing a community to select for a desired microbial function. We assume here that we have a library of variants with different quantitative traits, and our selection scheme favors variants with the best detoxification properties.

**Fig 1.**
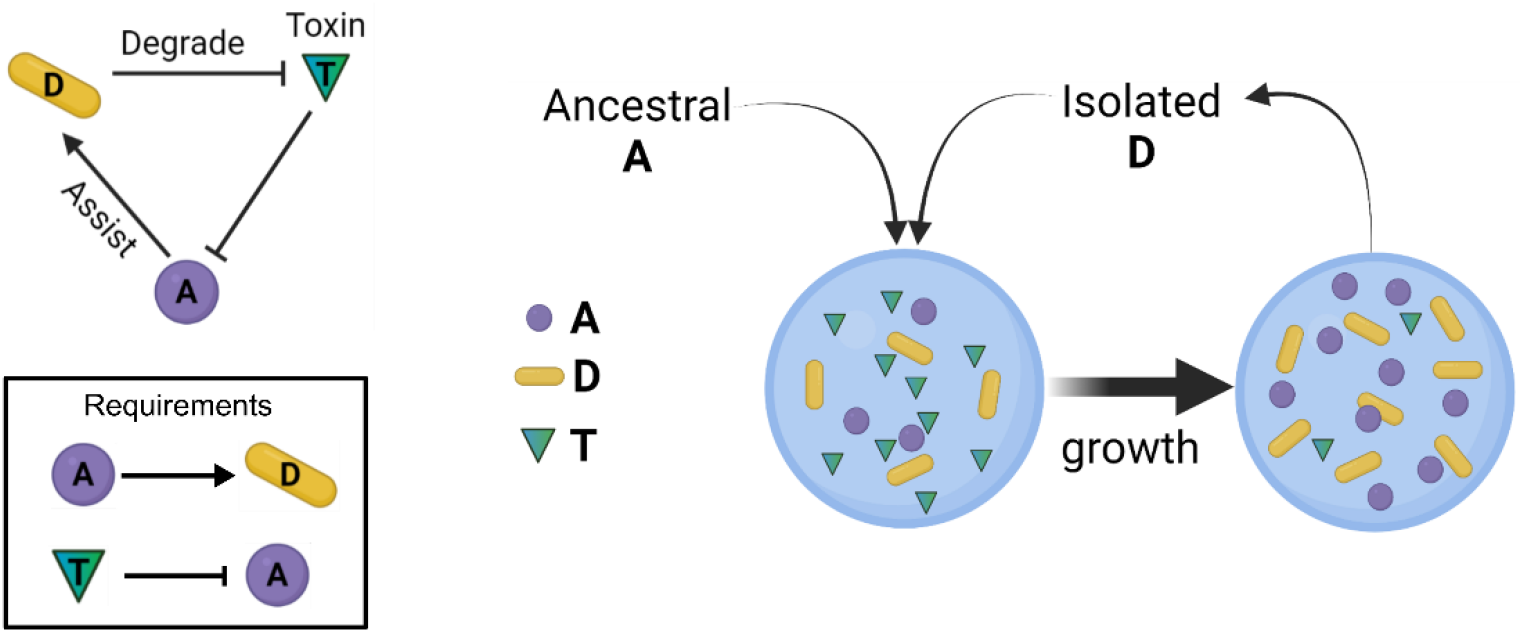
An assisting population A can generate a positive feedback for D from the toxin T. The overall scheme and the specific requirements are shown on the left. On the right, a conceptual selection scheme is illustrated in which cycles of coculture (with ancestral **A** and evolved **D**) leads to improved detoxification performance of **D**. We envision a droplet-based implementation where **D** is clonal within each culture but different droplets contain different variants of **D**.

## Results

### Introducing an assisting population can generate fitness feedback for degraders

The situation we envision is when a species **D** is identified that can degrade a specific toxin **T**, but the toxin has no fitness impact, positive or negative, on **D**. To allow selection for improved degradation activity by **D**, we propose to introduce an assisting population, **A**, with two requirements (Fig 1, left): (1) **A** provides a growth benefit to **D**, and (2) **A** is sensitive to **T** and gets inhibited by it. Here we assume the direct interaction between **A** and **D** to be commensalism, with little direct impact on **A** from **D**. In a coculture of **A** and **D**, better degradation of **T** by **D** relieves the inhibition on **A** and in turn provides a positive influence on **D** through **A**. This positive feedback can be used then to select for variants of **D** that better degrade **T**. In our proposed selection scheme (Fig 1, right), in each round ancestral **A** will be paired with evolved **D** from a previous round to ensure that the evolutionary pressure is focused on **D**.

### An implicit model captures major aspects of population dynamics

To assess how well the scheme in Fig 1 will work, we employ an implicit model in which the impacts of **A** on **D, D** on **T**, and **T** on **A** are implicitly included as fitness contributions in a population-level model (Methods-Model 1). We will refer to this implicit model of interactions as ImpInt in what follows. As an example, Fig 2 shows the simulated dynamics of cell populations and toxin concentration starting from a given initial condition. In this example, population A grows at a slow rate initially (under inhibition by **T**) until its density is high enough to support the growth of population **D**. Rapid growth of population **D** leads to a rapid decrease in toxin **T** concentration, and in turn, a lower inhibition of population **A**. In this example, within ∼48 hours the toxin is completely depleted, before populations **A** and **D** reach their saturation levels.

**Fig 2.**
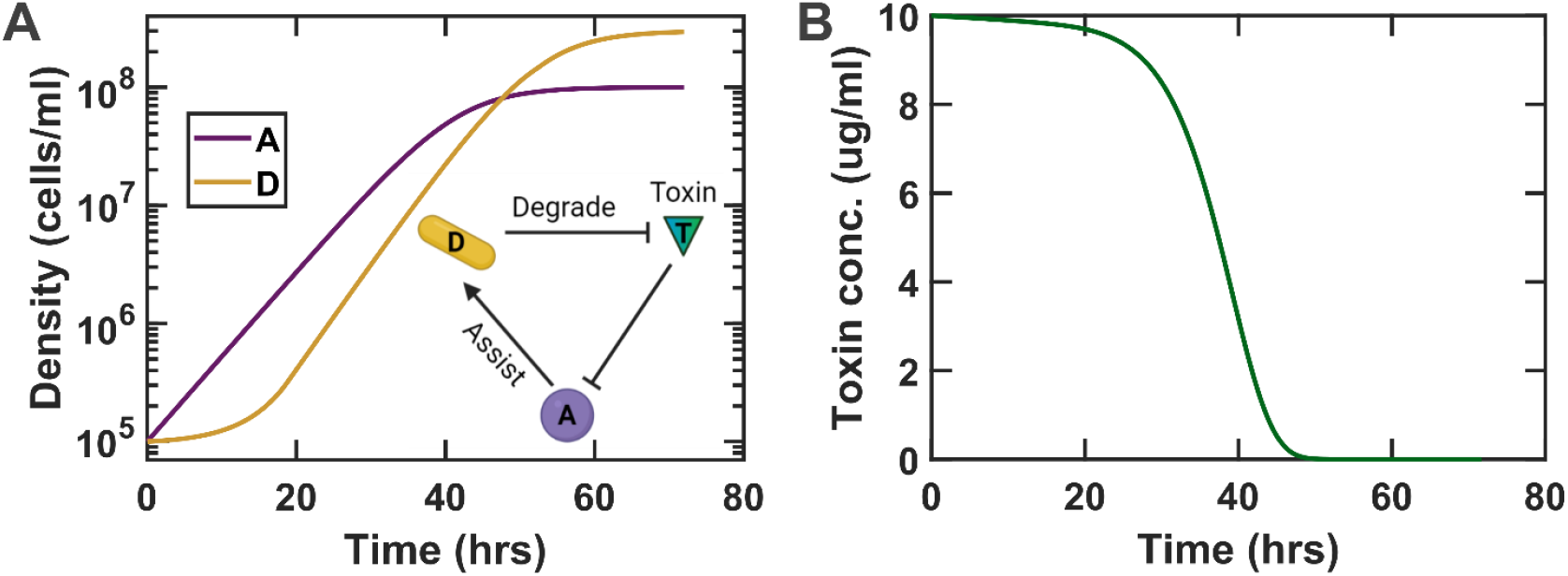
The assisting and degrading populations can grow together and degrade the toxin of interest. The dynamics of population densities (A) and the toxin concentration (B) are shown after incorporating all interactions. In the example shown here, populations **A** and **D** are assumed to be initially at 10^5^ cells/ml and the initial toxin concentration is 10 μg/ml. All relevant parameters are listed in Table 1. The ImpInt model is used for this simulation.

To assess whether the ImpInt model is adequate for representing this system, we explored two more explicit models: ExpEnz explicitly incorporates the **T**-degrading enzyme produced by **D** (Methods-Model 2), whereas ExpRes explicitly incorporates the resource produced by **A** that supports the growth of **D** (Methods-Model 3). We find that ImpInt can adequately approximate the dynamics of more explicit ExpEnz and ExpRes models (Figs S2 and S3). Given the agreement between the implicit models and the more explicit models, for simplicity we will use ImpInt in the remainder of our simulations. We note that one exception was when a strong enzyme decay rate was assumed in ExpEnz. For such a situation, a modified implicit model ImpLD (Methods-Model 4) had to be adopted, in which only growing **D** cells contribute to toxin degradation (Fig S4). The same approach used here can be used with ImpLD as well.

**Table 1.** Typical parameter values for the model.

| Parameter | Description | Value |
| --- | --- | --- |
| $t_f$ | Total simulation time per round | 60 hr |
| $r_A$ | Maximum growth rate of population **A** | 0.2 hr<sup>-1</sup> |
| $r_D$ | Maximum growth rate of population **D** | 0.22 hr<sup>-1</sup> |
| $K_A$ | Maximum carrying capacity of population **A** | 10<sup>8</sup> cells/ml |
| $K_D$ | Maximum carrying capacity of population **D** | 3×10<sup>8</sup> cells/ml |
| $\rho_T$ | Inhibition coefficient of **T** against **A** | 0.003 ml/(μg·hr) |
| $s_A$ | Growth coefficient of **A** in supporting **D** | $10^{-7}$ ml/(cells·hr) |
| $d_D$ | Detoxification coefficient of **T** removal by **D** | $10^{-8}$ ml/(cells·hr) |
| $d_E$ | Detoxification coefficient of **T** removal by **E** | $10^{-8}$ ml/(U·hr) |
| $\eta_D$ | Production rate of enzyme **E** by **D** | $2.5 \times 10^{-6}$ μU/(cells·ml) |
| $\beta_R$ | Production rate of resource **R** by **A** | 0.2 fmole/(cells·hr) |
| $K_R$ | Monod coefficient for growth of **D** on **R** | 0.2 μM |
| $\alpha_D$ | Consumption rate of resource **R** by **D** | 0.07 fmole/cell |
| $\delta_E$ | Decay rate of enzyme **E** | 0.02-0.5 hr<sup>-1</sup> |

### Geometric mean of A and D population sizes determines culture viability

We first asked under what condition a coculture of **A** and **D** is viable. It is expected that higher initial densities of **A** and **D** have higher propensity to be viable. We surveyed a range of initial densities of **A** and **D** in the model, which confirmed this expectation (Fig 3A). Additionally, it appeared from this survey that a higher initial density of **A** or **D** can compensate for a lower initial density of its partner. Probing further and replotting the same data as a function of the geometric mean of the initial densities of **A** and **D** (i.e. 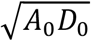), we observed that 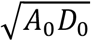 is a good predictor of whether a coculture is viable and how well it degrades the toxin within a given time (Fig 3B).

**Fig 3.**
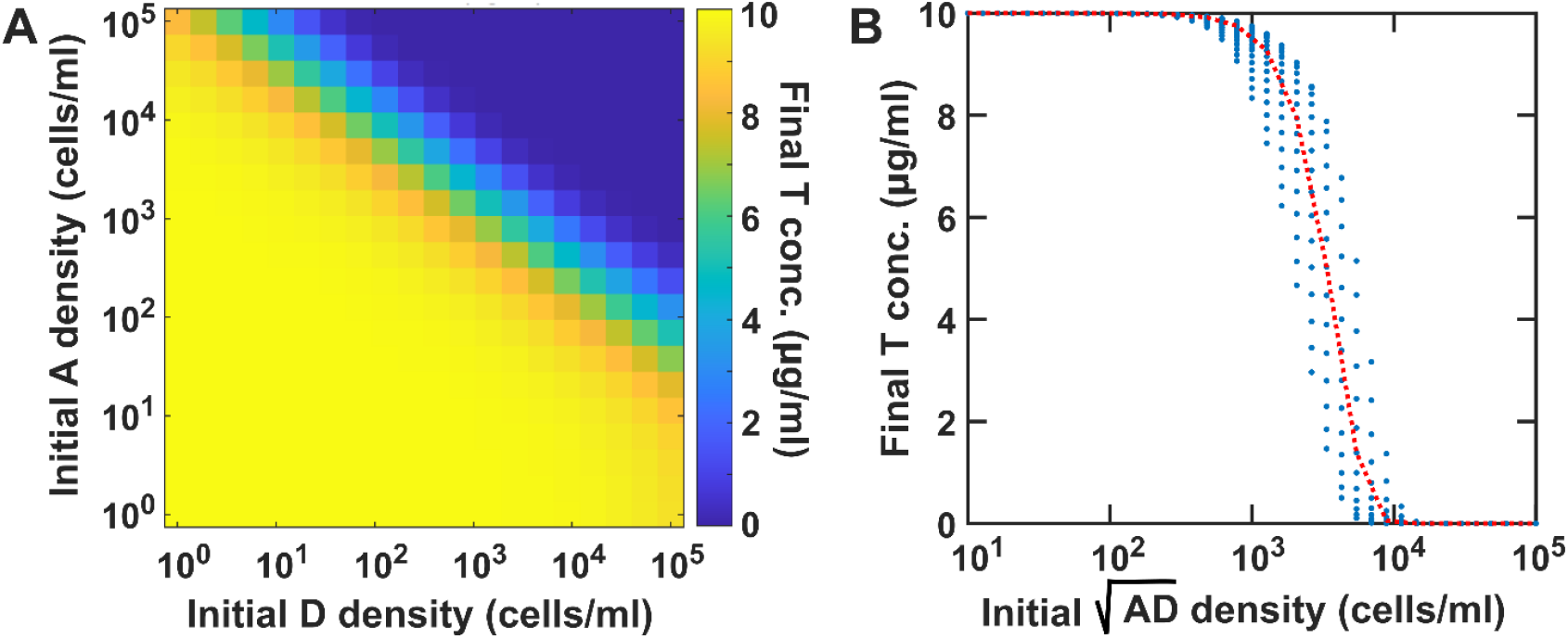
Viability of A-D cocultures depend on the geometric mean of the initial A and D population densities. (A) Surveying a range of initial **A** and **D** population densities shows that an increase in the initial density of one can compensate for a drop in the initial density of the other one to maintain viability. (B) Examining the final **T** concentrations suggest that the geometric mean of the initial **A** and **D** population densities is the main determinant of viability and degradation performance. Final **T** concentrations are taken from the simulations at 72 hours. In all cases the initial toxin concentration is 10 μg/ml. All relevant parameters are listed in Table 1. The ImpInt model is used in these simulations.

### Despite other sources of stochasticity, selection based on growth leads to improved detoxification

The main premise of our proposed PAAS scheme is that effective detoxification will be translated into improved overall culture growth—a trait that can be readily selected on. To assess the efficacy of such an approach, we computationally examined whether variants with better detoxification rates would be selected using PAAS. To create a more realistic situation, we assumed that in addition to the detoxification coefficient, other properties of the population (including their growth rates, carrying capacities, inhibition coefficient of **A** by **T**, and growth support coefficient of **D** by **A**) also varied stochastically (see Table 2). We then simulated many conditions (n=10000 instances) with random assignments of these variables and examined the traits in the output.

**Table 2.** Different random variables and their distributions in a typical artificial selection simulation.

| Random variable | Distribution | Value |
| --- | --- | --- |
| $r_A$ | Skew-normal | $\mu = r_A; \sigma = 0.02r_A; \text{skew } \alpha = -3$ |
| $r_D$ | Skew-normal | $\mu = r_D; \sigma = 0.02r_D; \text{skew } \alpha = -3$ |
| $K_A$ | Normal | $\mu = K_A; \sigma = 0.02K_A$ |
| $K_D$ | Normal | $\mu = K_D; \sigma = 0.02K_D$ |
| $\rho_T$ | Normal | $\mu = \rho_T; \sigma = 0.02\rho_T$ |
| $s_A$ | Normal | $\mu = s_A; \sigma = 0.02s_A$ |
| $d_D$ | Normal | $\mu = d_D; \sigma = 0.2d_D$ |

First, we found that the detoxification rate (*d*_*D*_) showed a positive association with overall growth, measured in total cell density (Fig 4A). Additionally, the overall detoxification performance was correlated with the total cell density, as expected (Fig 4B). To examine the efficacy of selection, we compared the distributions of the detoxification rates before selection and after selecting the top 10% instances with the highest total cell densities. This selection in PAAS clearly exhibits a preference for higher detoxification rates (Fig 4C). These results confirm the capability of PAAS to select for improved detoxifiers. Additionally, PAAS offers the advantage that growth as the primary trait of interest is relatively easy to measure, compared to direct measurements of the toxin concentration, e.g. through fluorescence [13], ELISA, or HPLC [14,15].

**Fig 4.**
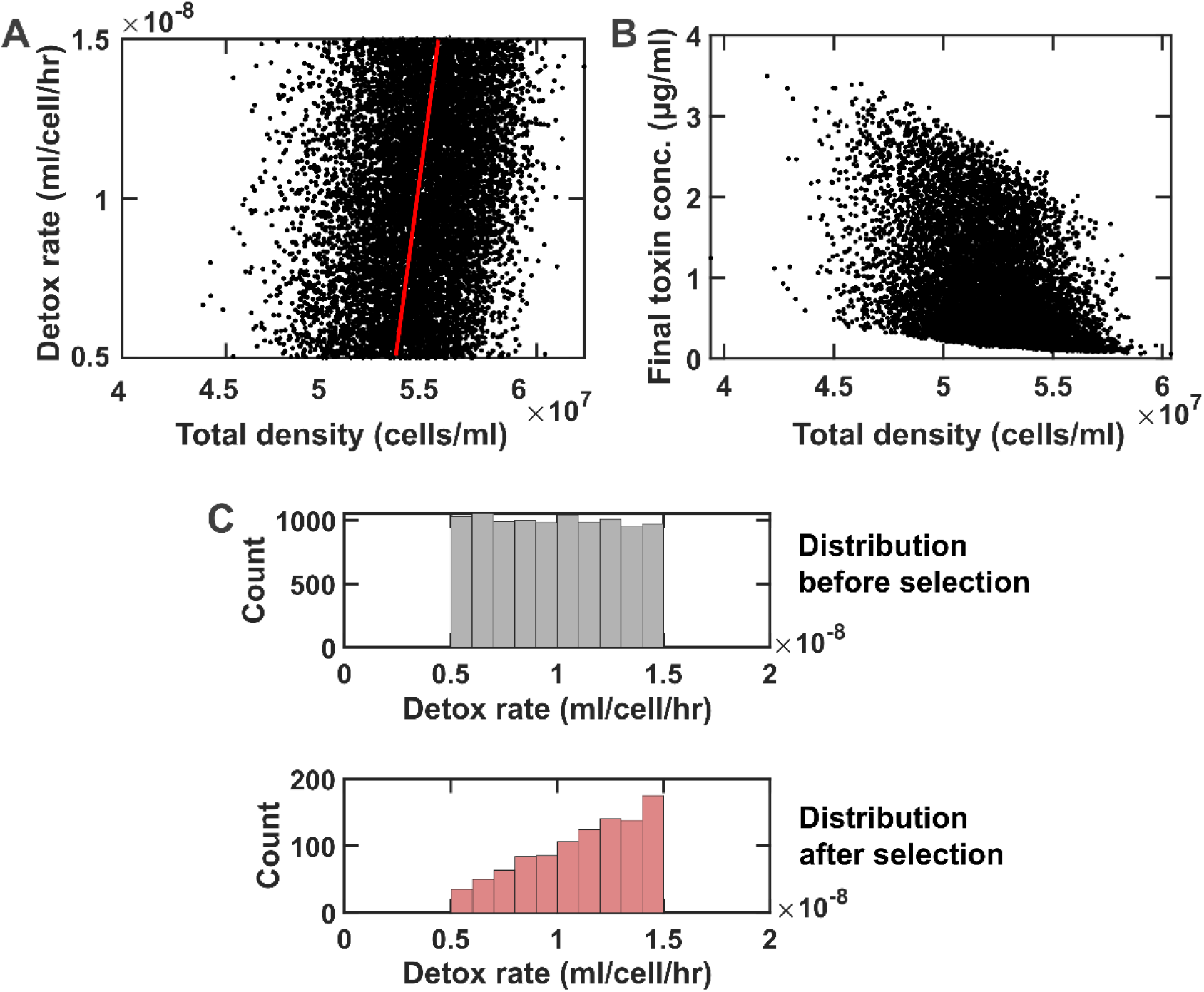
A survey of many (n=10000) simulated instances with stochastic parameters shows that PAAS allows us to select for improved detoxification as a secondary function. (A) Scatter-plot of all instances shows a positive correlation between the detoxification rate and total cell density. The red trend line is estimated based on the average total cell densities at low and high detox rates. (B) Total cell density is also tightly linked to the effectiveness of detoxification. (C) Comparing the distributions of the detoxification rates before selection and after selecting the top 10% instances with the highest total cell densities shows that PAAS favors improved detoxification. Final **T** concentrations are taken from the simulations at 44 hours. In all cases the initial toxin concentration is 10 μg/ml. All relevant parameters are listed in Table 1 and stochastic properties are listed in Table 2. The ImpInt model is used in these simulations.

### Effective detoxification selection is sensitive to the timing of propagation

To assess the efficacy of the selection scheme, we used detox improvement as a measure of improvement in function, defined as the average detoxification rate of selected instances compared to that of initial instances. We first assessed how the initial composition of the coculture affected detox improvement. Interestingly, the selection performance—as estimated by detox improvement—was higher in a particular range of initial densities (Fig 5A). Further investigation revealed that this range corresponded to initial cell densities that resulted in **T** being mostly, but not completely, degraded. In fact, examining the data based on the residual **T** after 60 hours of simulated growth showed a clear trend with detox improvement being maximum around 1% residual **T** and dropping to lowed values when residual **T** was much higher or lower (Fig 5B). This trend is intuitively expected; with too little or too much degradation, there is little information for resolving which cultures are performing well for degradation. We additionally examined the effect of the time between inoculation and passage and the results, consistent with the effect of initial density (Fig 5), that low, but not too low, residual **T** leads to the best detox improvement (Fig S5).

**Fig 5.**
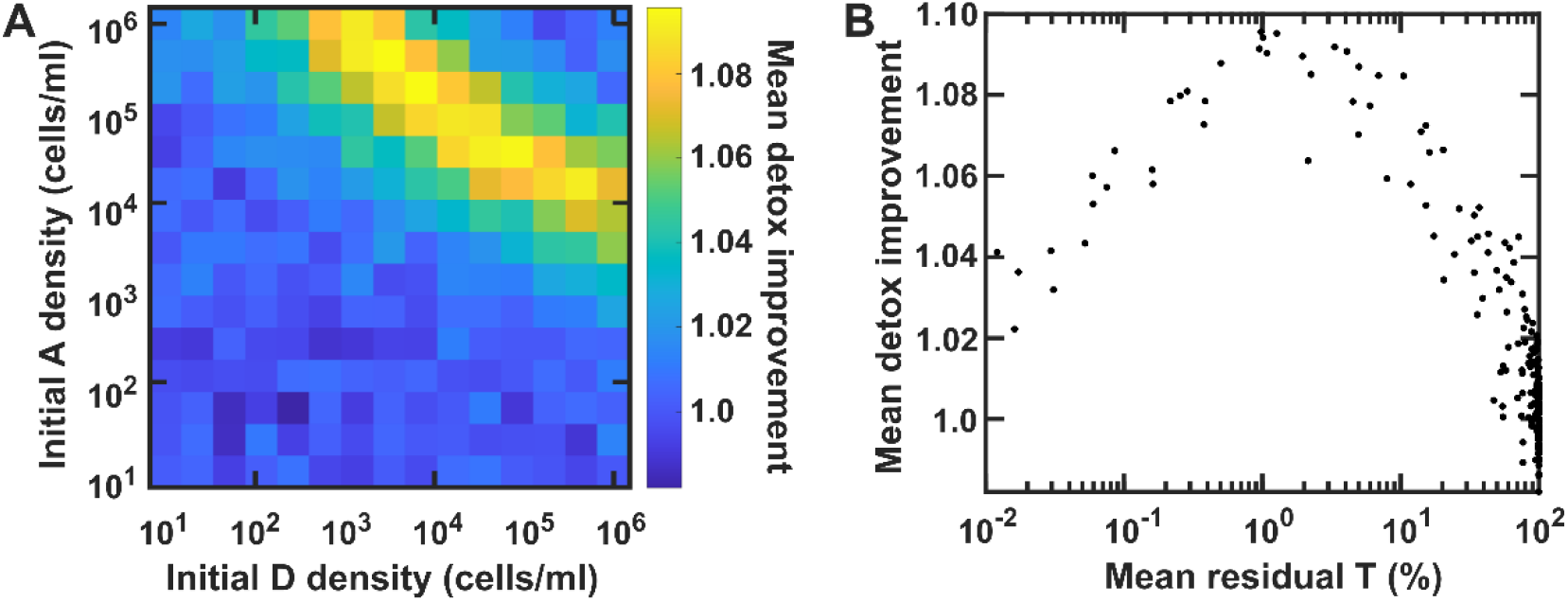
For optimal selection, most, but not all, of the toxin should be degraded at the time of selection. (A) Mean detox improvement (defined as the average of detoxification coefficients at the end of a round divided by its initial value) is plotted as a function of initial population densities of **A** and **D**. (B) Mean detox improvement data in (A) is plotted as a function of the final residual **T**, showing an optimal performance around 1% residual **T** at the end of each round. For each data point, 1000 instances were sampled, with stochastic parameters listed in Table 2. Final **T** concentrations are taken from the simulations at 60 hours. In all cases the initial toxin concentration is 10 μg/ml. All relevant parameters are listed in Tables 1 and 2. The ImpInt model is used in these simulations.

### Detoxification selection depends on the population bottleneck

Selection is expected to depend on the size of the bottleneck. With a more stringent bottleneck (i.e. selecting more extreme cases), the expectation is to get more extreme phenotypes, but at the risk of added uncertainty of losing the best performers. We asked if the same considerations applied to the PAAS scheme. We constructed 100 samples of the PAAS scheme, with *n* = 100 instances of coculture (with stochastic parameters as in Table 2) in each of the samples. For each of these cases, we enforced a range of bottlenecks, from choosing the top 1% total cell density, to choosing the top 30%. The results showed that, as expected, the outcome of less stringent bottlenecks was more consistent, but on average led to lower improvement (Fig 6A). Defining *bottleneck stringency* as the fraction of the total number of instances to the instances selected, we saw a saturable improvement with more stringent bottlenecks (Fig 6B). Importantly, the uncertainty in detox improvement was directly related to how stringent the bottleneck was, with 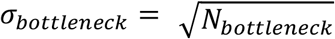, and *N*_*bottleneck*_ as the size of the selected instances in the bottleneck (Fig 6C). Overall, these trends follow the expectations for a standard selection scheme.

**Fig 6.**
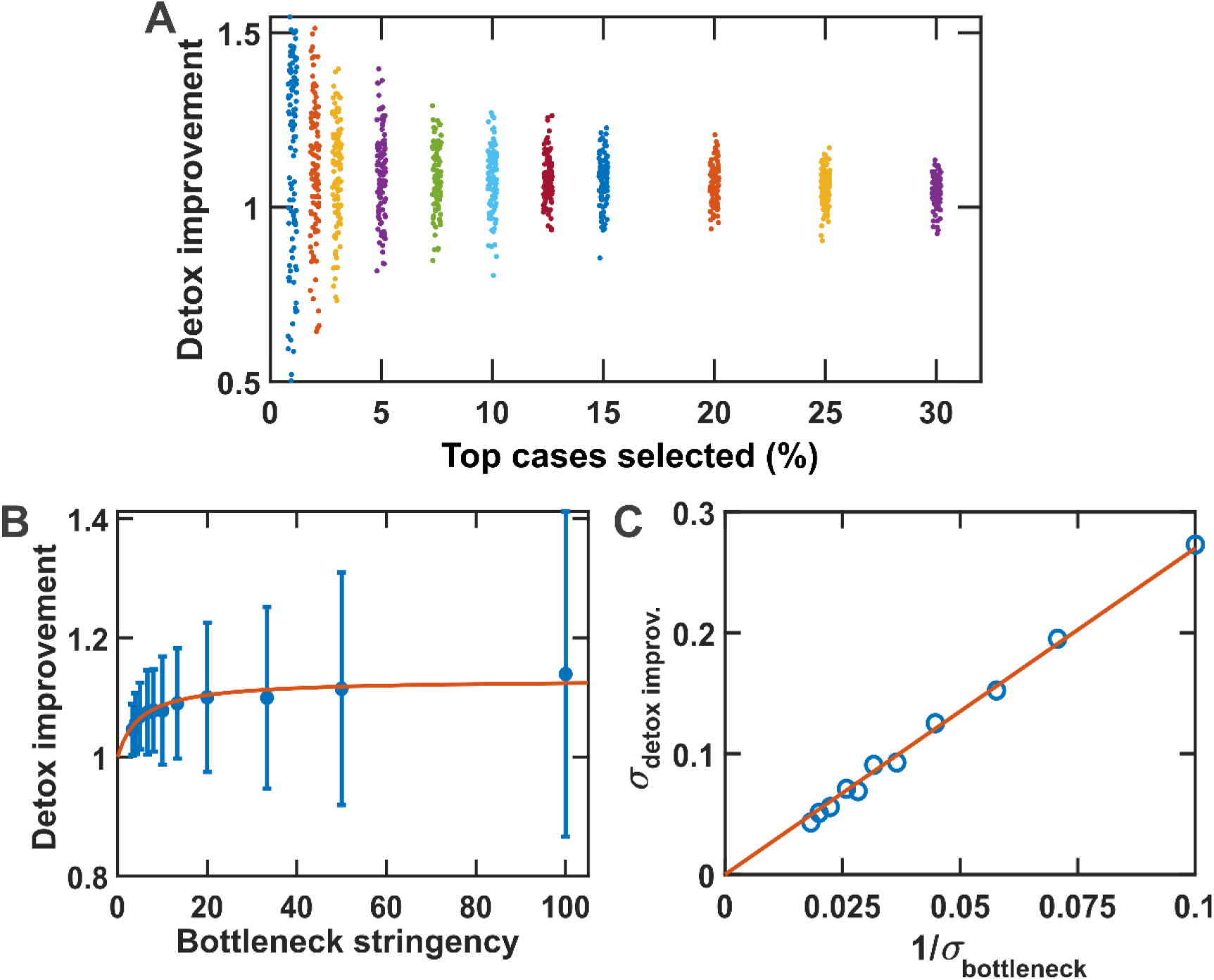
Improvement in detox, as a function of population bottleneck. (A) The distribution of detox improvement values are shown when different fractions of the top cases with the most growth are selected within a round. More stringent selections can potentially yield higher detox improvement, but at the risk of more uncertainty. (B) Error bars are standard deviations (n = 100). Red curve is a fit into the data, with the form *y* = 1 + (*y*_*f*_ − 1) *x*/(*x* + *x*_*s*_), where *y*_*f*_ = 1.3 and *x*_*s*_ = 5. (C) Red curve is a linear fit into the data, *y* = *mx*, where *m* = 2.7. Final **T** concentrations are taken from the simulations at 72 hours. In all cases the initial toxin concentration is 10 μg/ml. All relevant parameters are listed in Table 1. The ImpInt model is used in these simulations.

### Stochasticity in other cell traits can disrupt effective selection in PAAS

Stochasticity in other parameters is one of the main factors that can potentially derail the PAAS selection scheme by muddying which cultures are the best detoxifiers. We examined how different parameters correlated with the total cell density as our main selection criterion (Fig S6). We then asked how much stochasticity in other parameters can be tolerated in PAAS. For this, we examined a range of different values of standard deviations for the parameters listed in Table 2. We found that excessive stochasticity in other traits could mask the degradation performance of the cocultures (Fig 7). This was evident as the correlation between detoxification coefficient and the total cell density (i.e. our criterion for selection) is lost when stochasticity in other traits is large (Fig 7A). As a result, our selection for improved detoxification is no longer effective in such cases (Fig 7B).

**Fig 7.**
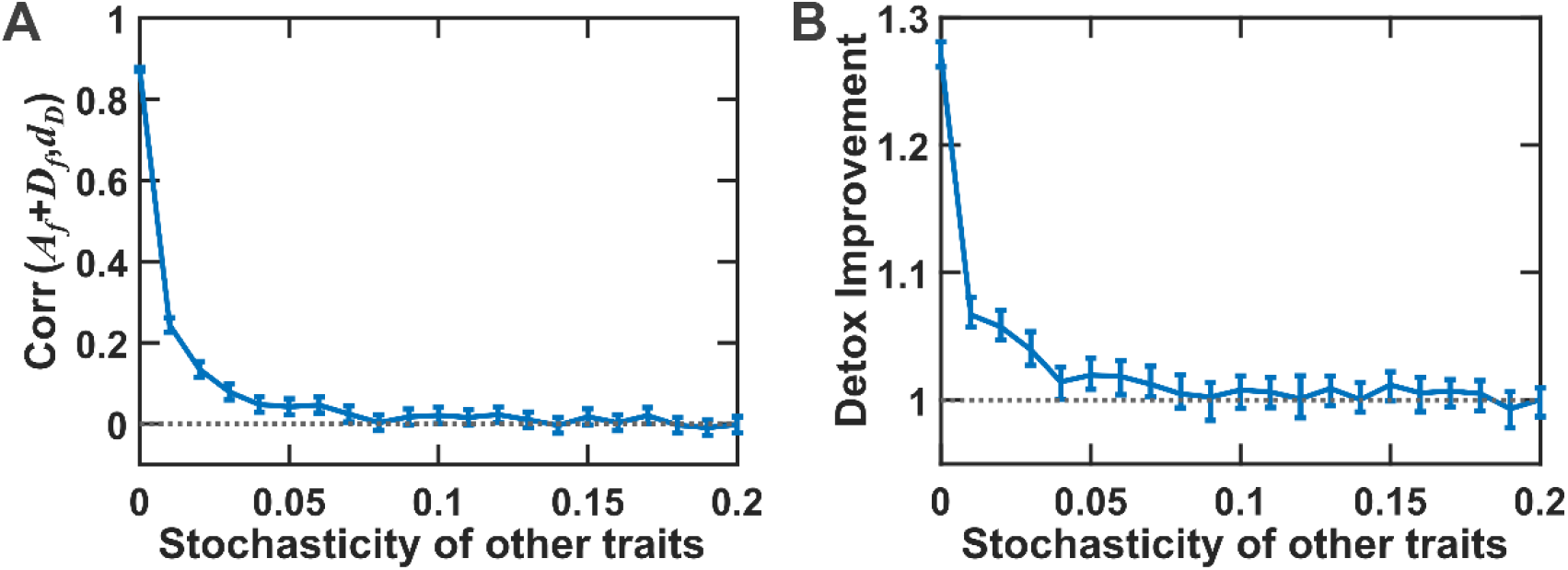
Stochasticity in other traits can interfere with PAAS efficiency. (A) Correlation between total cell density and detoxification coefficient decreases as stochasticity in other traits increases. Correlation coefficient is calculated using all instances of cocultures with parameters picked from corresponding random variables. Stochasticity is defined as the ratio of σ to µ (see Table 2) and the same value is used for all random variables except **d**_**D**_ for which σ/µ is fixed at 0.2. Error bars are standard deviations calculated using the top 10% of instances selected based on total cell density. (B) Detox improvement decreases with more stochasticity in other traits. Here, error bars depict bootstrap 95% confidence intervals using 100 samples of PAAS. Top 10% of the instances with the largest total population densities are selected for calculating detox improvement. All relevant parameters are similar to Fig 4 and listed in Tables 1. The ImpInt model is used in these simulations.

## Discussion

We investigated the capabilities and limitations of a partner-assisted artificial selection scheme to select for functions of interest that have no significant fitness impact on the cells that provide them. We introduced an assisting population that created a feedback between the function of interest and the fitness of the function provider. To investigate the potentials and limits of PAAS, we examined a system consisting of a toxin degrader, along with an assisting population that was sensitive to the toxin of interest and beneficial to the degrader population. We found that selection for overall population density can lead to improved detoxification rates. This selection is most effective if selection happens when detoxification is close to complete, so that there is enough discrimination between degraders with different performance. We see that bottleneck considerations in PAAS largely mirror our expectations in standard selection schemes. A more stringent bottleneck leads to a saturating improvement in detoxification performance, but at the cost of more uncertainty. Finally, we observe that too much stochasticity in other traits can mask the performance of toxin degradation and interfere with PAAS selection.

For practical implementation, we note that initial population sizes and the timing of selection can be used as effective design parameters. One major decision for designing an effective PAAS is the choice of bottleneck stringency; our *in silico* model suggests that PAAS is similar to a standard selection scheme in terms of how a more stringent bottleneck leads to stronger, but more uncertain, selection. Another major decision is the treatment of other sources of stochasticity. Among stochastic parameters that could interfere with selection, the growth rates of **A** and **D** appear to play major roles (Fig S6). Since the **A** population is reintroduced at the beginning of each round (Fig 1, right), a pre-adaptation step to maximize its growth rate can significantly reduce the variability in this trait. In contrast, the growth rate of **D**, as long as it does not come at the cost of loss of degradation capabilities, could be considered a desired trait to select for.

In our treatment of different traits, we have assumed that such traits are independent of each other. However, some correlation between these traits is possible, for example a positive or negative correlation between the growth rate and carrying capacity of cells [16]. If known, such correlations can be directly incorporated into the model for a more realistic representation of stochasticity.

Another assumption in our model is that **A** and **D** engage in commensalism and there is little direct impact on **A** by **D**, be it positive or negative. This can be controlled to some extent by choosing **A** that satisfies this assumption. We expect results similar to the condition examined in this manuscript with weakly positive or negative impact on **A** by **D**. Strong positive or negative impact on **A** by **D** can turn this commensalism into mutualism or exploitation regimes, respectively. Extreme exploitation conditions could drive **A** out of the community and disrupt PAAS. In contrast, strong mutualism is expected to stabilize the population dynamics [17] and lead to a more balanced performance regardless of the initial conditions.

Overall, we propose that PAAS can be utilized as an additional tool to expand the power of selection to situations where the function of interest has little fitness influence on the provider of that function. We recognize that an actual implementation will likely involve adjusting the scheme to the specifics of a system of interest. Our simplified model presented in this work offers a baseline to build upon.

## Materials and Methods

### Bacterial growth characterization

*Rhodococcus erythropolis* (DSM 43066) was grown from the frozen stock in glucose-yeast-malt (GYM) at 28° C with continuous shaking (240 rpm) for 24 hrs before starting the experiments. For the growth characterization experiment, *R. erythropolis* was cultured in basal Z medium: KH_2_PO_4_ (1.5 g/L), K_2_HPO_4_ × 3H_2_O (3.8 g/L), (NH_4_)_2_SO_4_ (1.3 g/L), sodium citrate dihydrate (3.0g/L), FeSO_4_ (1.1 mg/L), glucose (4.0 g/L), 100x vitamin solution (1 mL), 1000x trace elements solution (1 mL), 1 M MgCl_2_ (5 mL), 1 M CaCl_2_ (1 mL), and 100x amino acid stock (10 mL). AFG_2_ stock (Cayman Chemical) was dissolved in LC-MS grade methanol to the final concentration of 1 mg/mL. AFG_2_ was then introduced into the growth cultures at different concentrations by further diluting the stock in methanol to keep the total methanol concentration fixed across all cases.

Final volumes of 150 µl per well were used in standard flat-bottom 96-well plates. A BioTek Synergy Mx multi-mode microplate reader was used to monitor optical density of cells at 600 nm. Reads were taken at 5 min intervals over 48 hrs. Cultures usually started at an initial OD of 0.01 and were continuously shaking between reads. Five replicates were used per condition. Only the internal wells of the 96-well plate were used for samples, and the peripheral wells of the plate were filled with sterile water to contain evaporation.

Growth rates were calculated using a Matlab code that extracted the data from text files generated by BioTek Synergy Mx. The function `‘fit_logistic’` (written by James Condor, and available at https://www.mathworks.com/matlabcentral/fileexchange/41781-fit_logistic-t-q) was used to estimate the growth rates from OD readings.

### Models and equations

There are three assumptions shared in our models. (1) The growth rate of assisting population **A** linearly decreases as the concentration of the toxin **T** increases [18]. (2) The growth rate of **A** and its carrying capacity proportionally change at different concentrations of an inhibitor [19]. (3) Degradation rate of **T** is proportional to the population density of the degrading population **D**. Among these assumptions, only the second assumption is necessary for the overall PAAS scheme described in this manuscript. Nonetheless, we have included these assumptions to make the models more realistic, while keeping them simple.

#### Model 1: Implicit interaction effects (ImpInt)

In this simplified model, we assume logistic growth for the **A** and **D** populations. The toxin **T** is assumed to modulate both the growth rate and the carrying capacity of the population **A**. Growth rate and carrying capacity of population **D** is capped by the benefits supplied by population **A**.

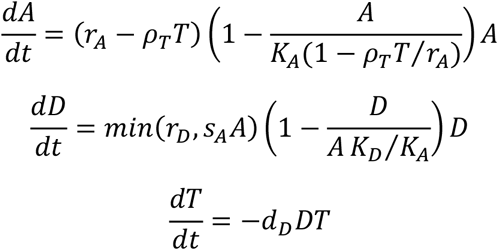

#### Model 2: Explicit enzyme effect (ExpEnz)

In this model, the **T**-degrading enzyme (produced by **D**) is explicitly included. Compared to ImpInt, rather than direct degradation of **T** by **D, D** produces the enzyme E which degrades **T**. We have also included an explicit term for intrinsic enzyme decay in our equations.

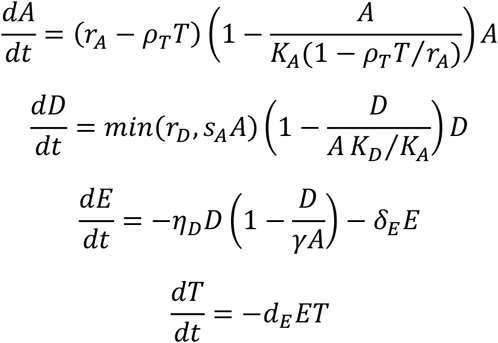

#### Model 3: Explicit resource effect (ExpRes)

In this model, the resource **R**, produced by **A** and supporting the growth of **D**, is explicitly included. We assume a standard Monod-type growth for D on R as its main limiting resource. The consumption of **R** by **D** is also assumed to be proportional to the biomass generated by the growing **D** population.

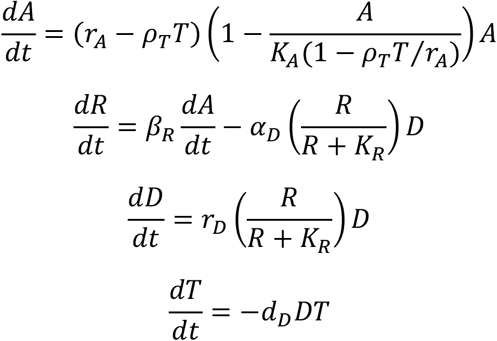

#### Model 4: Implicit interaction effects, live degradation (ImpLD)

In this modified phenomenological model, we assume that only growing **D** populations contribute to the detoxification. This will capture cases where the enzyme decay is large and thus detoxification stops when there is no growth and enzyme production by **D** cells.

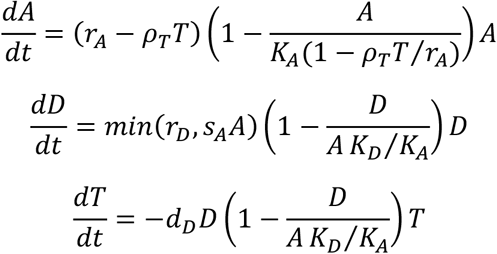

### Simulations

Numerical simulations were performed using MATLAB. Source codes along with descriptions of parameters are available at https://github.com/bmomeni/partner-assisted-artificial-selection.

### Parameters and their values

Unless otherwise stated, Table 1 lists the values of parameters used in our simulations. The order-of-magnitude of values are inferred from experimental characterization of aflatoxin G2 detoxification by *Rhodococcus* species.

### Random variables and statistics

Table 2 lists the distributions used for different random variables used to include stochasticity in our simulations. For all normal random variables, we used the built-in `random` function in Matlab, with relevant parameters (e.g. `‘uniform’` for a uniform distribution and `‘normal’` for a normal distribution). To generate skew normal distributions for growth rates, we used the following transformation based on two independent random variables *x*_1_ and *x*_2_ picked from a Normal distribution 𝒩(0,1).

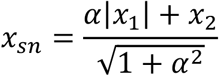

Here *α* is the skew parameter in the distribution. The distribution is more skewed towards small (/large) values, when *α* is negative (/positive).

Bootstrap confidence intervals are calculated using the `bootci` function in Matlab, with `mean` as the target function.

## Acknowledgments

BM and MZ are supported by the National Science Foundation (NSF-CBET) under Grant No. 2103545. NS is supported by a NIFA-AFRI Predoctoral Fellowship from the USDA (Award No. 2021-67034-35108). Figure 1 was created with <u>BioRender.com</u>.

## Supplementary Information

**Fig S1.**
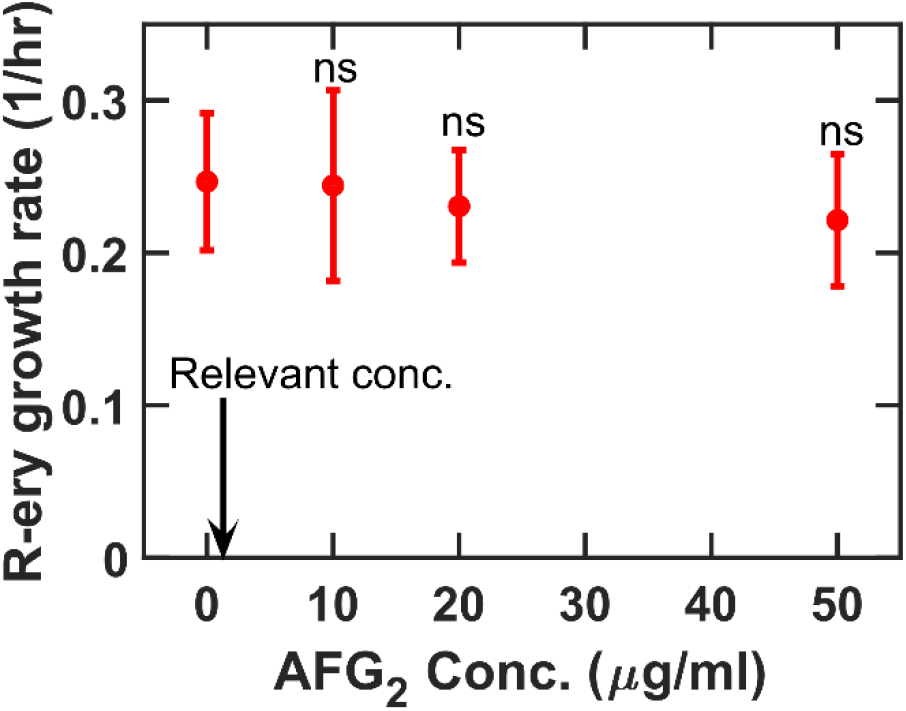
Growth rate of detoxifying strains such as *Rhodococcus erythropolis* is minimally affected by the presence of aflatoxins, highlighting the challenge of natural selection for improved detoxification. Different concentrations of AFG_2_ (dissolved in methanol) are added to basal Z culture medium (see Materials and Methods, Bacterial growth characterization) inoculated with *R. erythropolis* at an initial cell OD of 0.01. Cultures are allowed to grow and the initial growth rate of *R. erythropolis* is estimated from the increase in OD over time (as a proxy for cell density). None of the growth rates at 10, 20, or 50 μg/ml of AFG_2_ were statistically different from the no-toxin control (t test, p>0.3). For comparison, The upper limit of practically relevant concentrations of AFG_2_ (around 1 μg/ml) is marked by an arrow as a point of reference to show even at much higher AFG_2_ concentrations the fitness impact is minimal.

**Fig S2.**
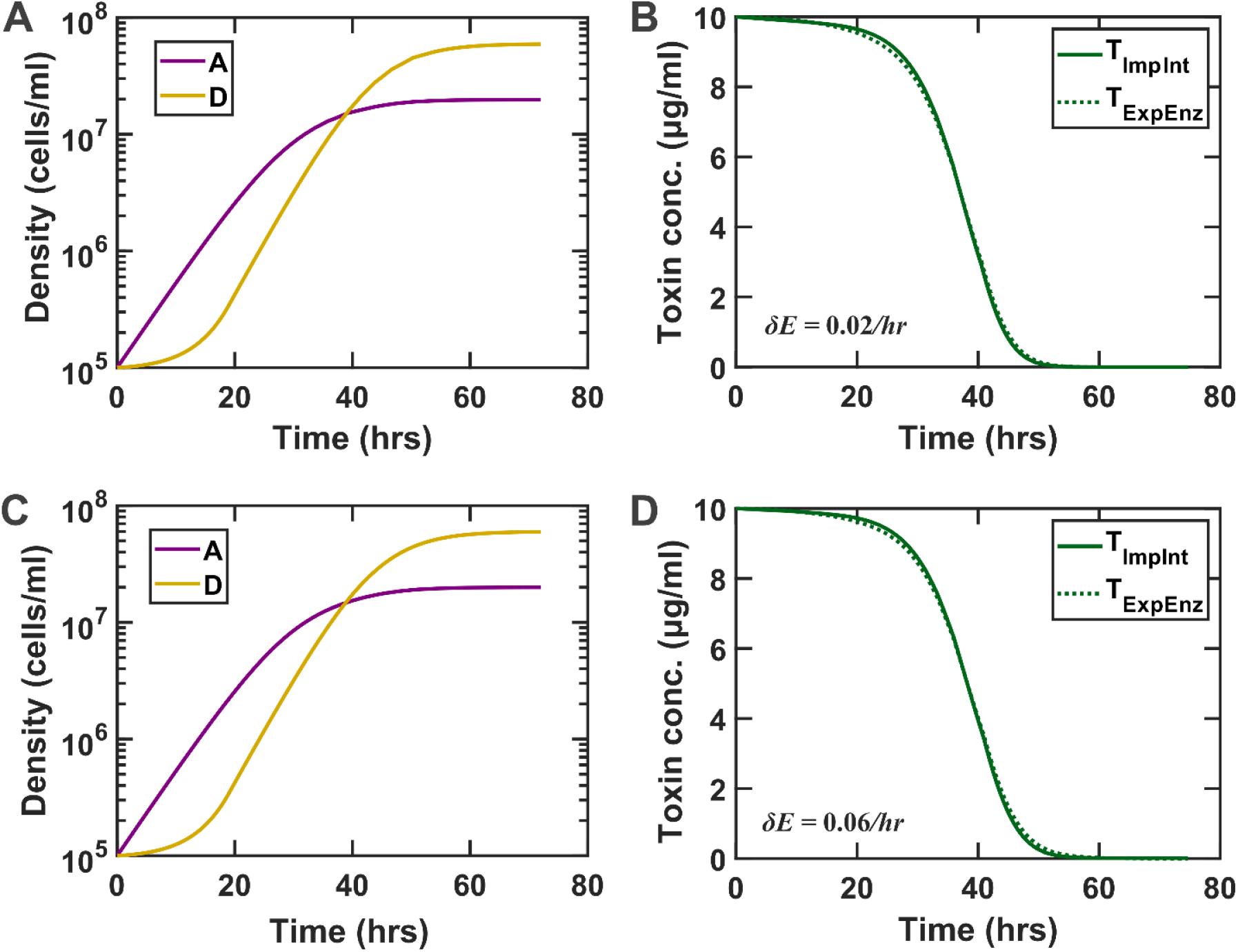
The simplified ImpInt model can adequately approximate a more mechanistic model that explicitly includes the degrading enzyme (ExpEnz). The equations behind ImpInt and ExpEnz models can be found in the Methods section (Model 1 and Model 2, respectively). The degradation coefficient in ImpInt is adjusted to match the dynamics of **T** offered by ExpEnz.

**Fig S3.**
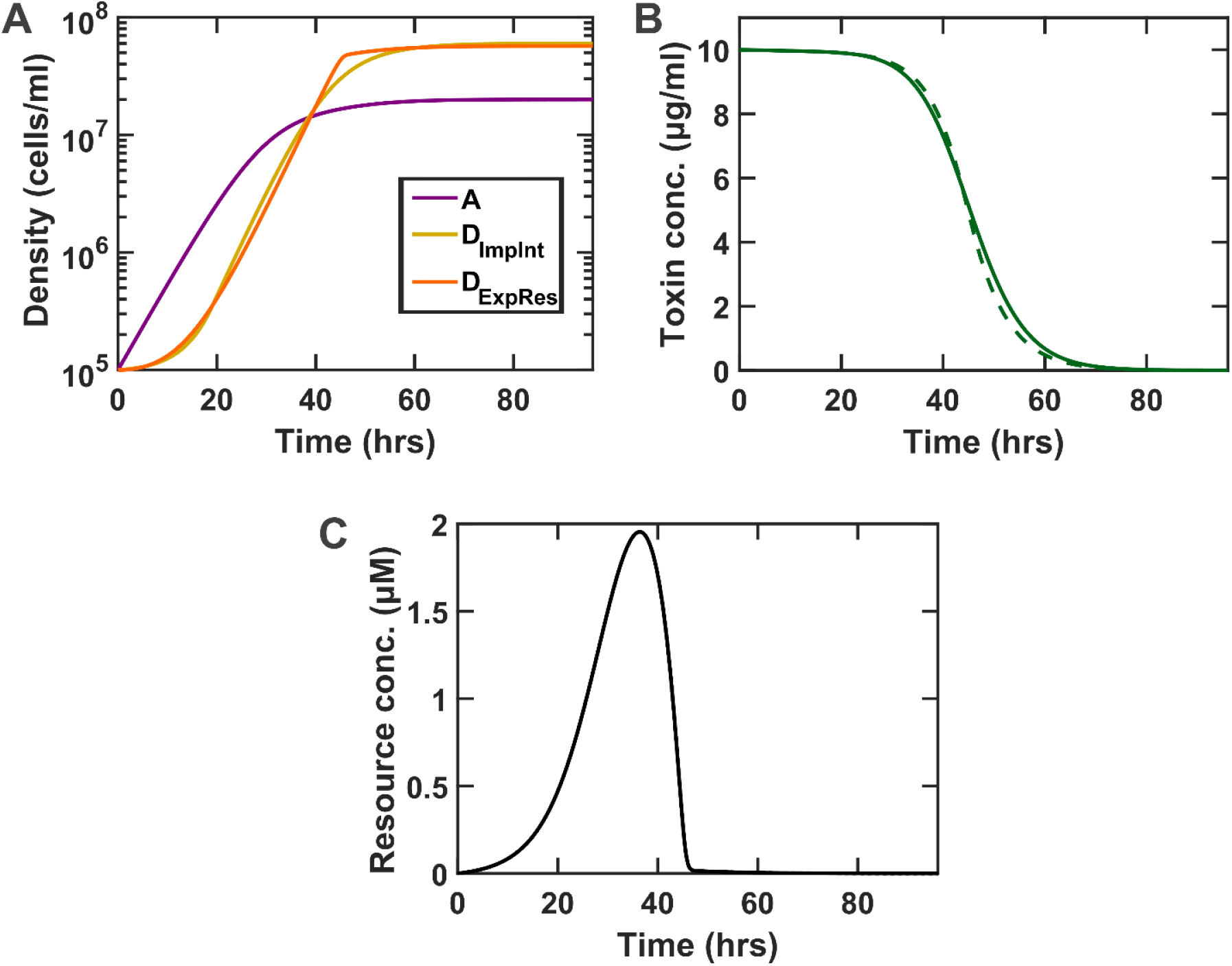
The simplified ImpInt model can adequately approximate a more mechanistic model that explicitly includes the resource or metabolite that mediates how population A supports population D (ExpRes). The equations behind ImpInt and ExpRes models can be found in the Methods section (Model 1 and Model 3, respectively). The degradation coefficient in ImpInt is adjusted to match the dynamics of **T** offered by ExpRes.

**Fig S4.**
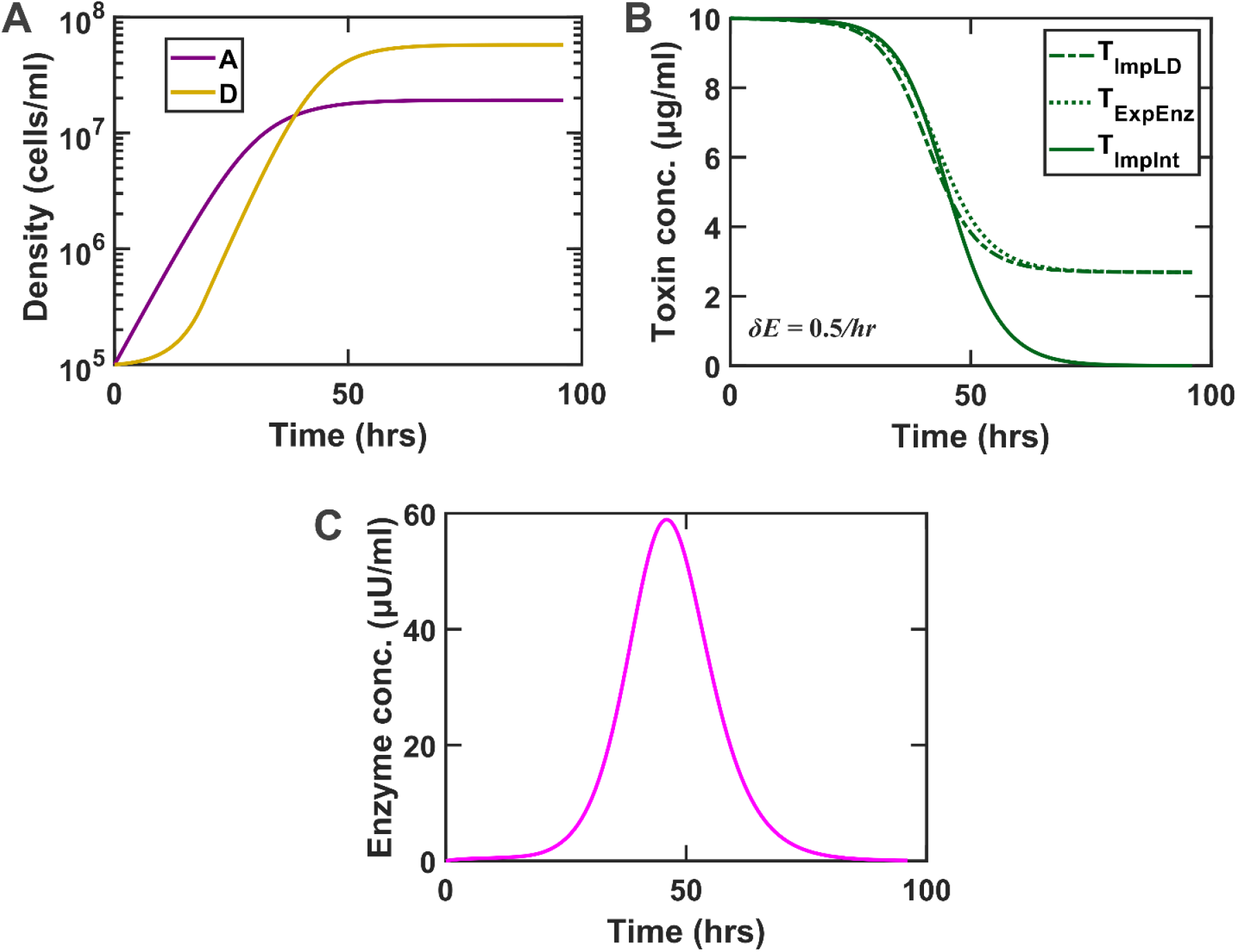
When enzyme decay rate is large, a modified implicit model that assumes degradation only by growing D cells (ImpLD) can adequately approximate the model that explicitly includes the degrading enzyme (ExpEnz). The equations behind ImpLD and ExpEnz models can be found in the Methods section (Model 4 and Model 2, respectively). The degradation coefficient in ImpLD is adjusted to match the dynamics of **T** offered by ExpEnz. We note that ImpInt no longer matches the dynamics of **T** from ExpEnz when the enzyme decay rate is very high.

**Fig S5.**
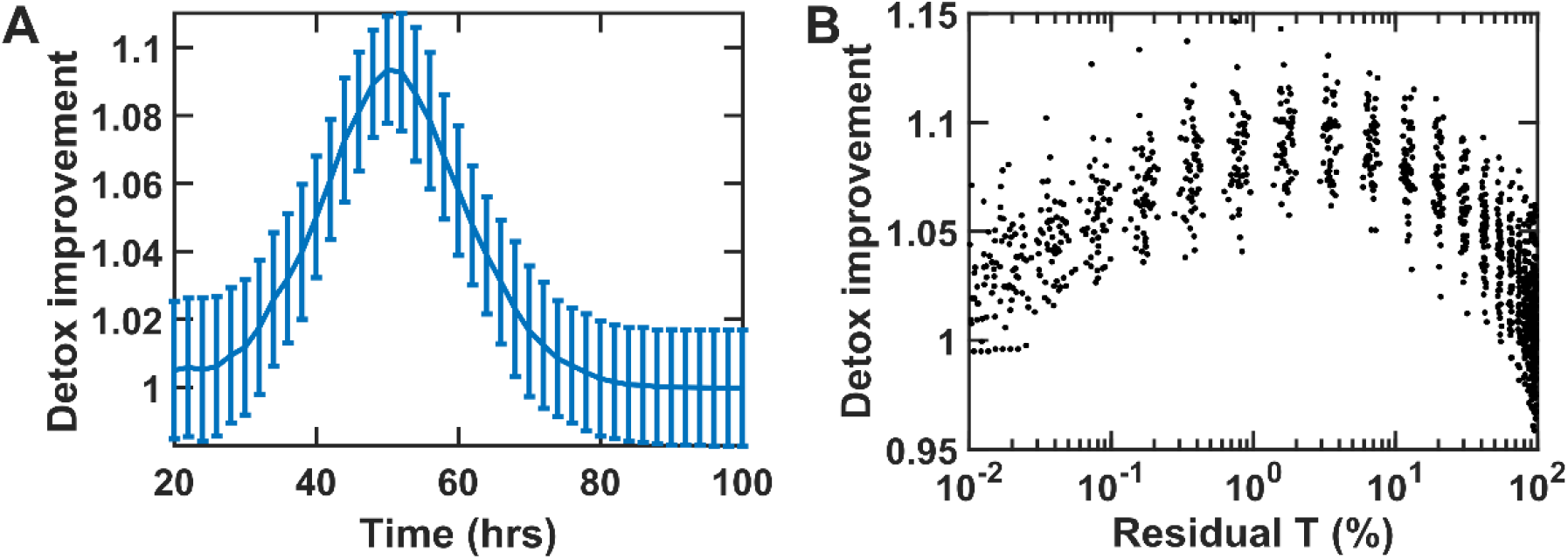
For optimal selection, most, but not all, of the toxin should be degraded at the time of selection. (A) Detox improvement (defined as the average of detoxification coefficients at the end of a round divided by its initial value) is plotted as a function of detoxification time. Error bars show standard deviations calculated among 50 independent instances. (B) Detox improvement data in (A) is plotted as a function of the final residual T, showing an optimal performance around 1% residual **T** at the end of each round. For each data point, 1000 instances were sampled, with stochastic parameters listed in Table 2. Initial **A** and **D** densities are 10^5^ cells/ml each. In all cases the initial toxin concentration is 10 μg/ml. All relevant parameters are listed in Tables 1 and 2, except *K*_*A*_ = 2×10^7^ cells/ml and *K*_*D*_ = 6×10^7^ cells/ml. The ImpInt model is used in these simulations. All the parameters match those in Fig 5.

**Fig S6.**
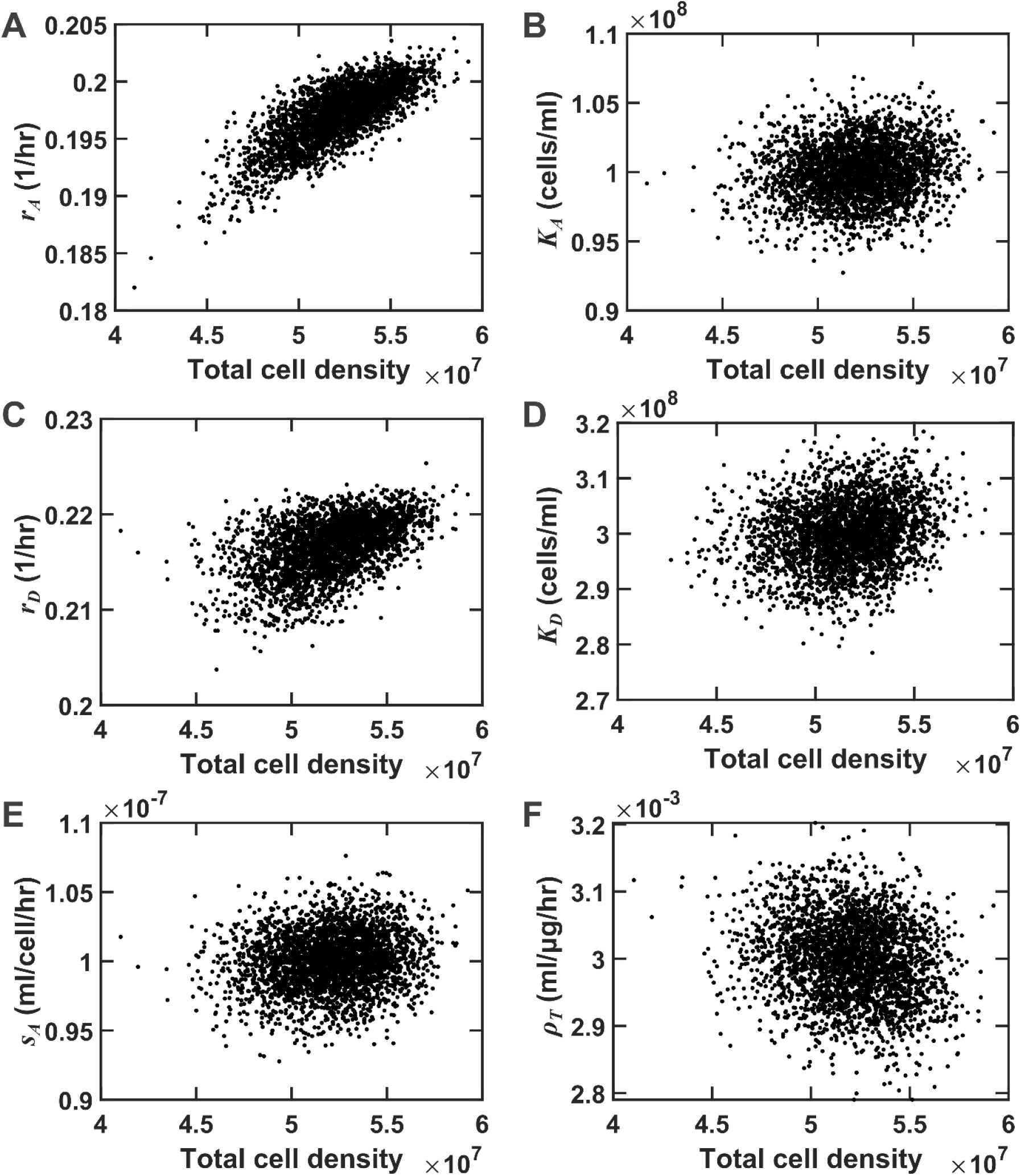
Stochasticity in growth rates of A and D as the major contributors to the total cell density can interfere with detoxification selection. We survey n=3000 simulated instances with stochastic parameters to evaluate how stochasticity in parameters affects PAAS selection. (A-F) Scatter-plots show that among different parameters, *r*_*A*_ and *r*_*D*_ are the most influential in determining the total cell, and can thus interfere with our ability to select for improved detoxification. Total cell density is found from simulations at 72 hours. The initial toxin concentration is 10 μg/ml. All relevant parameters are listed in Table 1 and stochastic properties are listed in Table 2. The ImpInt model is used in these simulations.

